# SHARPIN serine 146 phosphorylation mediates ARP2/3 interaction, cancer cell invasion and metastasis

**DOI:** 10.1101/2022.01.21.477220

**Authors:** Umar Butt, Meraj H Khan, Jeroen Pouwels, Jukka Westermarck

## Abstract

The adaptor protein SHARPIN is involved in a number of cellular processes and promotes cancer progression and metastasis. However, how the choice between different functions of SHARPIN is post-translationally regulated is unclear. Here we have characterized SHARPIN phosphorylation by mass spectrometry and *in vit*ro kinase assay. Focusing on two uncharacterized phosphorylation sites, serine 131 and 146, in the unstructured linker region of SHARPIN, we demonstrate their role in SHARPIN-ARP2/3 complex interaction, whereas they play no role in integrin inhibition or LUBAC activation. Consistent with its novel role in ARP2/3 regulation, serine 146 (S146) phosphorylation of SHARPIN promoted lamellopodia formation. Notably, CRISPR-Cas9 mediated knockout of SHARPIN abrogated three-dimensional (3D) invasion of several cancer cell lines. The 3D invasion of cancer cells was rescued by overexpression of the wild-type SHARPIN, but not by SHARPIN S146A mutant, identifying S146 as an invasion promoting phosphorylation switch. Finally, we demonstrate that inhibition of phosphorylation at S146 significantly reduces the *in vivo* metastasis in the zebrafish model. Collectively, these results demonstrate that SHARPIN S146 phosphorylation constitutes a single functional determinant of cancer cell invasion both *in vitro* and *in vivo*.

## Introduction

The primary cause for cancer-related deaths is metastasis (Steeg, 2016). Significant improvements in cancer survival rates have been seen recently due to early diagnosis and development of targeted therapies (Guan, 2015). Metastasis however, remains still a hurdle that most cancer therapies are not able to overcome. Cancer metastasis involves several critical steps: first, the cancer cell(s) needs to detach from the primary tumor. Subsequentially, the detached cell needs to migrate into and through the surrounding tissue, a step called invasion. Then the metastasizing cancer cell needs to travel through the blood or lymph system, after which it needs to adhere to the secondary site, where it once more needs to invade to reach its final destination (Fares, Fares et al., 2020). Suppressing cancer metastasis by targeting any of these processes would be of an urgent therapeutic need(Ganesh & Massagué, 2021). However, this would require a detailed mechanistic understanding of how these processes are regulated, and consequently identification of potential target mechanisms for anti-metastatic therapies.

SHANK-associated RH domain interactor (SHARPIN) is mainly a cytoplasmic adaptor protein involved in the regulation of multiple cellular functions (De Franceschi, Peuhu et al., 2015, Gao, Bao et al., 2019, Jung, Kim et al., 2010, Khan, Salomaa et al., 2017, Liu, Wang et al., 2017, Park, Jin et al., 2016, Rantala, Pouwels et al., 2011, Zhang, Huang et al., 2014, Zhou, Liang et al., 2020). The most explored function of SHARPIN is its interaction with RBCK1 (HOIL) and RNF31 (HOIP) to form the Linear Ubiquitination Assembly Complex (LUBAC), a regulator of the canonical NF- κB pathway signalling (Gerlach, Cordier et al., 2011, Ikeda, Deribe et al., 2011, Tokunaga, Nakagawa et al., 2011). SHARPIN is also well known as an important inactivator of integrins (Pouwels, De Franceschi et al., 2013, Rantala et al., 2011). Other molecular targets of SHARPIN include T-cell receptor, caspase 1, EYA transcription factors, SHANK proteins, and PTEN (He, Ingram et al., 2010, Landgraf, Bollig et al., 2010, Lim, Sala et al., 2001, Nastase, Zeng-Brouwers et al., 2016, Park et al., 2016). Multiple cellular functions indicate that different signalling pathways compete for SHARPIN, and that SHARPIN functions as a signalling coordinator (De Franceschi et al., 2015). However, how differential binding of SHARPIN to its partners is spatio-temporally regulated remains unknown. Post-translational modifications (PTM) of SHARPIN are likely involved, as PTMs are known to function as molecular switches by affecting protein-protein interactions (Chen, Liu et al., 2020, Nishi, Hashimoto et al., 2011). However, besides the recent identification of S165 phosphorylation of SHARPIN as the activating phosphorylation for LUBAC activation (Thys, Trillet et al., 2021), the phosphorylation switches determining SHARPIN activity towards different cellular functions are as yet obscure.

SHARPIN gene is amplified, and at the protein level it is overexpressed in a variety of human cancers (Fig. S1A) (Bii, Rae et al., 2015, De Melo & Tang, 2015, He et al., 2010, Jung et al., 2010). The overexpressed SHARPIN promotes cancer cell proliferation, tumour formation and cancer metastasis (Bii et al., 2015, He et al., 2010, Li, Lai et al., 2015, Zhang et al., 2014). However, the molecular determinants by which these different cancer related functions of SHARPIN are regulated are poorly understood. What is known is that SHARPIN regulates cell adhesion and migration either by inhibition of integrins (De Franceschi et al., 2015, Pouwels et al., 2013, Rantala et al., 2011), or by promotion of lamellipodium formation through the ARP2/3 complex (Khan et al., 2017). The seven subunit ARP2/3 complex is responsible for creating branched actin networks through polymerization of actin (Blanchoin, et al.,2000, Rana, Alkrekshi et al., 2021). Overexpression of the ARP2/3 complex has been observed in a variety of human cancers (Iwaya, Oikawa et al., 2007, Liu, Yang et al., 2013, Otsubo, Iwaya et al., 2004, Semba, Iwaya et al., 2006, Zhang, Guan et al., 2012). This overexpression of the ARP2/3 complex is strongly associated with tumour cell invasion (Mondal, Di Martino et al., 2021), and can be used as a marker to differentiate benign lesions and malignant melanomas (Kashani-Sabet, Rangel et al., 2009). Therefore, understanding of mechanism that activates tumor invasion promoting ARP2/3 functions could lead to novel therapeutic opportunities for preventing metastasis, which is the dominant cause for death of cancer patients.

Here, we demonstrate that phosphorylation on SHARPIN at serine 146 promotes cancer cell invasion. This phosphorylation switch selectively mediates SHARPIN interaction with ARP2/3 complex indicating that this protein interaction might provide a target for therapeutic interference in cancer.

## MATERIALS AND METHODS

### Antibodies

These antibodies were used: rabbit Sharpin (14626-1-AP, Proteintech; 1:1000 WB), mouse cortactin (p80/85) (05-180, Merck Millipore; 1:300 IF), mouse GAPDH (5G4MaB6C5, HyTest; 1:20.000 WB), Alexa Fluor 488 Phalloidin (Invitrrogen; 1:300 IF)

These secondary antibodies were Alexa Fluor 488- or Alexa Fluor 555-conjugated IgGs (Invitrogen; IF), horseradish peroxidase (HRP)-conjugated IgGs (GE Healthcare; WB), DyLight 680- or 800-conjugated anti-mouse and rabbit IgGs (Thermo Scientific; WB). Mouse P5D2 (Hybridoma bank; 1:20 FACS), mouse 12G10 (ab30394, Abcam; 1:100 FACS)

### Plasmids and siRNAs

Construction of siRNA1-insensitive GFP–Sharpin and Sharpin mutant plasmids (De Franceschi et al., 2015) has been previously described. Construction of Arp3–TagRFP, (Khan et al., 2017) has been previously described. siRNAs: Sharpin [Hs_SHARPIN_1 HP siRNA (Qiagen)], and control siRNA [AllStars negative control siRNA (Qiagen)].

### Cells and transfections

HeLa cells were grown in Dulbecco’s modified Eagle’s medium (DMEM) with 10% fetal bovine serum (FBS), 1% L-glutamine, 1% MEM non-essential amino acids, 1% sodium pyruvate, 2% HEPES and 1% penicillin-streptomycin. HEK-293 cells were grown in DMEM with 1% penicillin-streptomycin, 10% FBS and 1% L-glutamine. NCI-H460 cells were grown in RPMI1640 with 10% FBS, 1% penicillin-streptomycin, 1% L-glutamine, 1% MEM non-essential amino acids, 1% sodium pyruvate and 1% glucose. Mda-Mb-231 cells were grown in DMEM with 10% FBS, 1% L-glutamine and 1% penicillin-streptomycin. PC3 cells were grown in RPMI with 10% FBS, 1% penicillin-streptomycin and 1% L-glutamine. All cell lines were regularly tested for contaminations and were from American Type Culture Collection (ATCC). Plasmid transfections were performed using Lipofectamine 2000 (HeLa and HEK-293 cells), Lipofectamine 3000 (NCI-H460 cells) (Life Technologies) and jetPRIME (MDA-MB-231 and PC3 cells). siRNA transfections were performed using Hiperfect (Qiagen).

### Recombinant proteins

Recombinant GST and GST–SHARPIN were produced in *E. coli* Rosetta BL21DE3 and purified according to the manufacturer’s instructions (BD Biosciences).

### IVK and Mass spectrometry

Recombinant kinases were purchased prom ProQinase GmbH. 20 ng of kinase was mixed with 1ug of GST-SHARPIN and incubated in 20 mM Hepes (pH 7.4), 10mM CaCl_2_, 25 mM MgCl_2_, 1 mM ATP and 5 μCi 32P-γ-ATP. Samples were then incubated on a heat block for 1 hour at 30 °C. Kinase reaction was terminated by using 2x Laemmli (SDS) sample buffer. Samples were then boiled at 100°C for 10 minutes and run on a gel. Comassie stained SDS-PAGE gel bands of GST-SHARPIN were then cut out for mass spectrometry. Protein samples were then digested by trypsin. Phosphopeptide enrichment was done by TiO_2_ chromatography. LC-MS/MS analysis was done using Q Exactive (a quadrupole-orbitrap mass spectrometer). Data analysis was done using Mascot database search against SwissProt E. coli supplemented with GST-tagged Sharpin and common contaminants.

For in cellulo analysis of SHARPIN phosphorylation GFP pulldowns were performed using GFP-Trap beads (ChromoTek) according to the manufacturer’s protocol. GFP-SHARPIN had been isolated by IP using beads and separated by SDS-PAGE. GFP-SHARPIN was in-gel digested by trypsin. Digested and desalted peptide samples were dissolved in 1% formic acid and analysed by LC-ESI-MS/MS using a QExactive mass spectrometer. The LC-ESI-MS/MS analyses were performed on a nanoflow HPLC system (Easy-nLC1000, Thermo Fisher Scientific) coupled to the QExactive mass spectrometer (Thermo Fisher Scientific, Bremen, Germany) equipped with a nano-electrospray ionization source. Peptides were first loaded on a trapping column and subsequently separated inline on a 15 cm C18 column (75 μm x 15 cm, ReproSil-Pur 5 μm 200 Å C18-AQ, Dr. Maisch HPLC GmbH, Ammerbuch-Entringen, Germany). The mobile phase consisted of water with 0.1% formic acid (solvent A) and acetonitrile/water (80:20 (v/v)) with 0.1% formic acid (solvent B). A linear 10 min gradient from 8% to 43% B was used to elute peptides. MS data was acquired automatically by using Thermo Xcalibur 3.0 software (Thermo Fisher Scientific). An information dependent acquisition method consisted of an Orbitrap MS survey scan of mass range 300-2000 m/z followed by HCD fragmentation for 10 most intense peptide ions.

The data files were searched for protein identification using Proteome Discoverer 1.4 software (Thermo Fisher Scientific) connected to an in-house server running the Mascot 2.4.1 software (Matrix Science) against SwissProt_2016_01 database. PhosphoRS 3.1 tool was used for detecting localization of phosphorylation sites.

### FACS

HeLa cella were seeded on to a 6 well plate. Next day cells were transfected with Control or Sharpin siRNA. Following day these cells were transfected with GFP control, GFP SHARPIN WT, GFP SHARPIN S131A or GFP SHARPIN S146A. The following day cells were harvested and fixed with 4% PFA. Cells were stained for active β1-integrin (12G10) or total β1-integrin (P5D2). Samples were analysed using FACSCalibur with CellQuest software (BD Biosciences) and non-commercial Flowing Software ver. 2.5 (Perttu Terho; Turku Centre for Biotechnology, Finland; www.flowingsoftware.com). The Integrin Activation Index was calculated by dividing the background-corrected active cell-surface integrin levels by total cell-surface integrin levels.

### NF-κB Reporter Assay

HeLa cella were seeded on to a 6 well plate. Following day these cells were transfected with Renilla Luciferase control vector (pRLTK), NF-κB reporter plasmid (pGL4.32[luc2P/NF-κB-RE/Hygro]) and WT or mutant GFP-SHARPIN expression plasmids. A GFP-only expression vector was used as a negative control. The next day, medium was replaced with medium with or without 50 ng/ml TNF, and after 5 h the luciferase activity was measured using the Dual-Luciferase Reporter Assay System (Promega), according to manufacturer’s instructions. Luminescence detection was done using Synergy H1 Multi-Mode Reader.

### FRET measurements by FLIM

HeLa cells were transfected with donor alone [GFP–Sharpin constructs (WT, or Phospho mutants) or with donor together with the acceptor (Arp3–TagRFP). Cells were fixed 24 hours post transfection and mounted with Mowiol 4-88 (Sigma-Aldrich). GFP fluorescence lifetime was measured by using a fluorescence lifetime imaging attachment (Lambert Instruments) on a Zeiss AXIO Observer D1 inverted microscope (Zeiss). For sample excitation, a sinusoidally modulated 3W, 497 nm LED at 40 MHz under epi-illumination was used. Cells were imaged using the 63× NA 1.4 oil objective (excitation, BP470/40; beam splitter, FT495; emission, BP525/50). The phase and modulation were determined using the manufacturer’s software from images acquired at 12 phase settings. Fluorescein at 0.01 mM, pH 9 was used as a lifetime reference standard. The FRET efficiency was calculated as previously described (Khan et al., 2017).

### Immunofluorescence

NCI-H460 were seeded in a 6 well plate. Next day cells were transfected with Control or Sharpin siRNA. Following day cells were trypsinized and re-seeded on to coverslips in a 24 well plate. On the following day the cells were transfected GFP control, GFP Sharpin WT, GFP SHARPIN S146A or GFP SHARPIN S146E and GFP SHARPIN V240A/L242A. Cells were fixed with 4% paraformaldehyde for 15 minutes at room temperature the following day. Permeabalization of cells was done with 0.1% Triton-X 100. Blocking was done with 10% goat serum. Cells were then stained with mouse cortactin (p80/85) overnight at 4 °C. Cells were imaged using Zeiss AxioVert 200 M inverted wide-field microscope equipped with a Plan-NEOFLUAR 63×1.25 NA oil objective (Zeiss) and Orca-ER camera (Hamamatsu Photonics). Image processing was performed using Fiji image analysis software (Schindelin, Arganda-Carreras et al., 2012).

### Sharpin-knockout cell lines created with CRISPR

Sharpin knockout NCI-H460 cell line was previously generated (Khan et al., 2017). Sharpin-knockout cell lines (MDA-MB-231, HeLa, PC3) were created using CRISPR genome engineering as previously described (Khan et al., 2017).

### Western Blotting

For assessing GFP-SHARPIN WT and mutans expression levels HeLa cells were seeded on to a 6 well plate. Next day the cells were transfected with GFP only, GFP SHARPIN-WT and mutant constructs. 48h post transfection cells were harvested, lysed run in SDS-PAGE. Proteins were transferred on to Nitrocellulose membranes and probed with anti-GFP and anti-GAPDH antibodies. For validation of SHARPIN CRISPR knockouts respective cell lines were grown on 6 well plates and harvested for western blots. Membranes were probed with anti-SHARPIN and anti-GAPDH antibodies.

### Cell proliferation

MDA-MB-231 SHARPIN CRISPR WT and KO cells were seeded on to a 96 well plate (1000 cells per well). Cells were then imaged every 2h using IncuCyte Zoom™ System (Essen BioScience) with a 10× objective for 4 days.

### Inverted invasion assay

Inverted invasion assays have been previously described (Jacquemet, Baghirov et al., 2016). Collagen 1 (concentration 5 μg ml−1; PureCol EZ Gel, Advanced BioMatrix) supplemented with Fibronectin (25 μg ml−1) was incubated at 1 hour at 37 °C to polymerize it in the inserts (8 μm ThinCert; Greiner bio-one). Inserts were then inverted and cells were seeded on the opposite side of the filter and allowed to attach to the matrix for 4h at 37 °C. The inserts were then place in serum free medium. Medium supplemented with 10% FBS was placed on top of the matrix in the inserts providing a serum gradient. Cells were fixed after 24-48 hours of seeding. 4% PFA was used to fix cells for 2 hours. Cell permeabilization was done using 0.5% Triton-X 100 at room temperature for 30 min. Cells were stained with Alexa-488 Phalloidin overnight at 4°C. Following staining the plugs were washed 3 times with PBS and imaged on a confocal microscope (LAM510; Ziess, LSM 780; Ziess, LSM 880; Ziess). Z-stacks of the samples were captured with a slice interval of 15um using a 20x objective lens (NA 0.50 air, Plan-neofluar). A montage of the individual confocal images is presented showing increasing penetrance from left to right. Invasion of cells was calculated using area calculator plugin in ImageJ. The fluorescence intensity of cells invading more than 45um was used to calculate the percentage of cells in the plug that were able to invade.

### Zebrafish embryo xenograft

Zebra fish were injected with MDA-MB 231 cells stably expressing either GFP-SHARPIN WT or GFP SHARPIN S146A mutant cells according to the previously described protocol (Paatero, Alve et al., 2018). Following transplantation embryos were imaged the next day using Zeiss SteREO Lumar.V12 microscope. After 4 days the embryos were imaged again. Image analysis was done using Fiji image analysis software (Schindelin et al., 2012). Cell populations representing distant metastasis were counted manually.

### Statistical analysis

All statistical analyses were performed using GraphPad Prism version 9 for Windows (GraphPad Software). The student’s t-test was used for normally distributed data (Shapiro-Wilk normality test α=0.05). For all other data, the Mann–Whitney test was used. P<0.05 was considered significant.

## Results

### Identification of *in vitro and in cellulo* SHARPIN phosphorylation sites

To better understand SHARPIN phosphorylation in cancer cells, an in vitro kinase assay (IVK) was performed with GST-SHARPIN in the presence of active forms of oncogenic kinases PKCalpha, CDK4/CycD3, FAK, ERK1, ERK2, AKT1 and AKT2 respectively. The autoradiograph revealed that GST-SHARPIN is phosphorylated by PKCalpha, CDK4/CycD3, ERK1 and ERK2 (Fig. 1A). Mass Spectrometry analysis of these samples revealed 12 phosphosites in GST-SHARPIN regulated by these kinases (Fig.1B)(Table 1).

**FIG 1.**
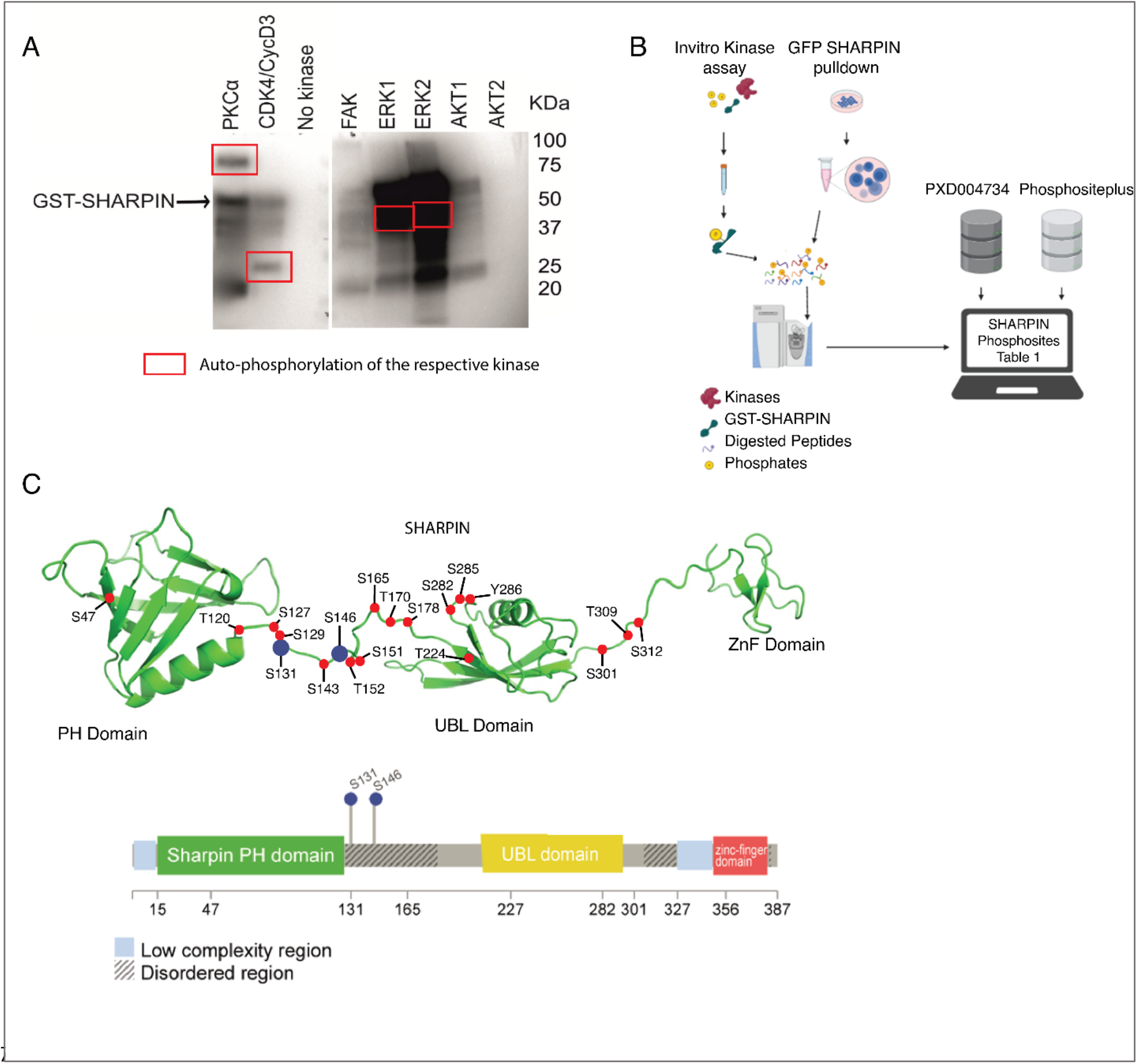
Phosphorylation of SHARPIN. (A) Sharpin phosphorylation by oncogenic kinases (PKCα, CDK4/CycD3, ERK1 and ERK2) in an In Vitro Kinase assay (IVK). (B) Schematic of approaches used for comprehensive analysis of SHARPIN phosphorylation. (C) SHARPIN phosphorylation sites on a cartoon model illustrating the individual functional domains connected by a linker region, and a lollipop diagram of SHARPIN showing phosphorylation sites in the disordered region selected for further analysis.

**Table 1.**
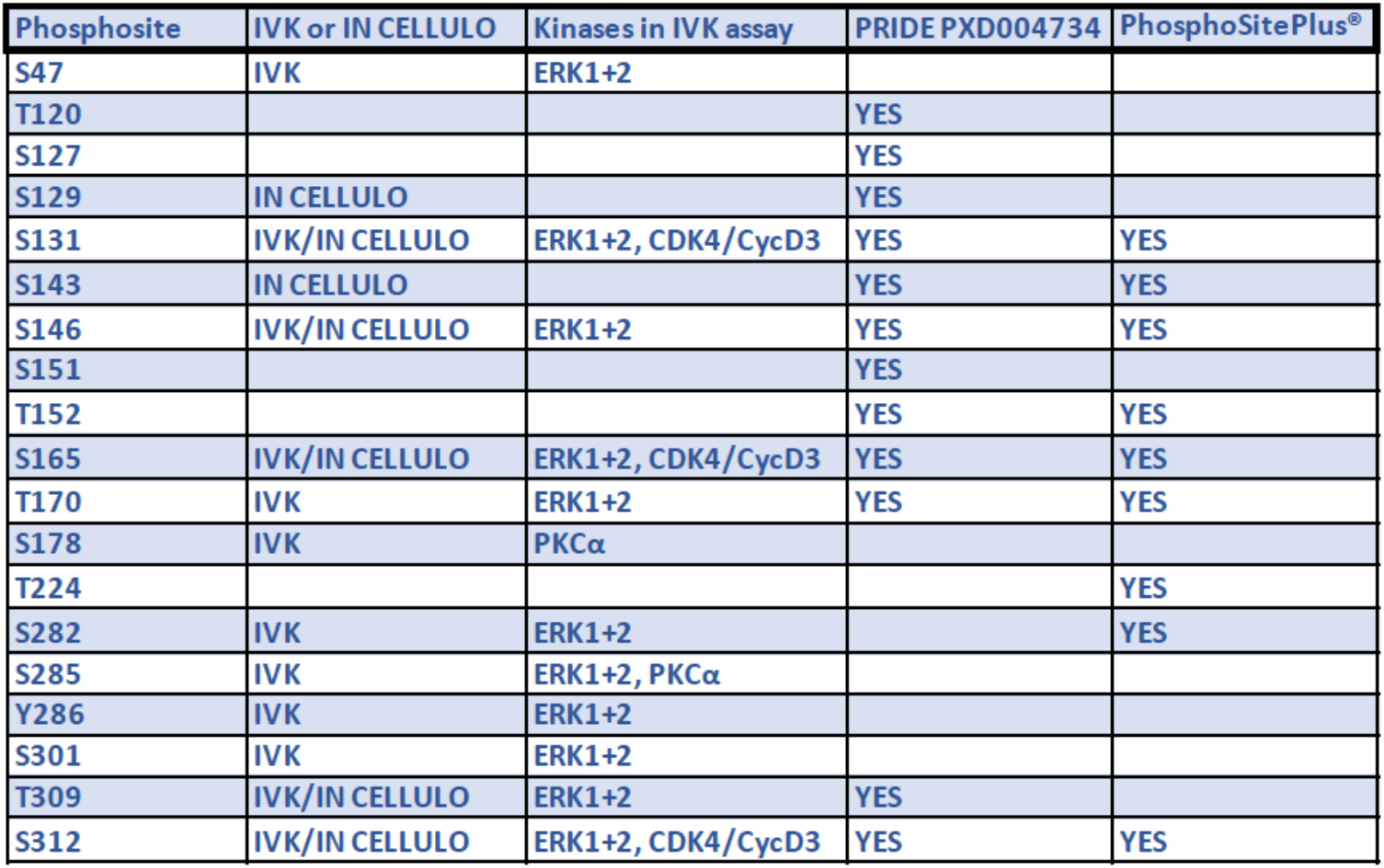
Summary of SHARPIN phosphorylation sites.

| Phosphosite | IVK or IN CELLULO | Kinases in IVK assay | PRIDE PXD004734 | PhosphoSitePlus® |
| --- | --- | --- | --- | --- |
| S47 | IVK | ERK1+2 |  |  |
| T120 |  |  | YES |  |
| S127 |  |  | YES |  |
| S129 | IN CELLULO |  | YES |  |
| S131 | IVK/IN CELLULO | ERK1+2, CDK4/CycD3 | YES | YES |
| S143 | IN CELLULO |  | YES | YES |
| S146 | IVK/IN CELLULO | ERK1+2 | YES | YES |
| S151 |  |  | YES |  |
| T152 |  |  | YES | YES |
| S165 | IVK/IN CELLULO | ERK1+2, CDK4/CycD3 | YES | YES |
| T170 | IVK | ERK1+2 | YES | YES |
| S178 | IVK | PKCα |  |  |
| T224 |  |  |  | YES |
| S282 | IVK | ERK1+2 |  | YES |
| S285 | IVK | ERK1+2, PKCα |  |  |
| Y286 | IVK | ERK1+2 |  |  |
| S301 | IVK | ERK1+2 |  |  |
| T309 | IVK/IN CELLULO | ERK1+2 | YES |  |
| S312 | IVK/IN CELLULO | ERK1+2, CDK4/CycD3 | YES | YES |

To identify which sites on SHARPIN are constitutively phosphorylated in proliferating cells, GFP pulldowns from 293T cells expressing GFP-SHARPIN or GFP alone were analyzed by affinity purification coupled with mass spectrometry (AP-MS)(Fig. 1B). The MS analysis revealed 7 SHARPIN phospho-sites; out of which serine 131 (S131), S146, S165, threonine 309 (T309), and S312 were overlapping with IVK sites (Table 1). Several phosphosites identified here had been observed also by a mass spectrometry analysis available in the Proteomics Identification Database (Identification of novel SHARPIN binders (PXD004734)) (Fig.1B)(Table 1). Moreover, 7 phosphosites of SHARPIN have been reported at https://www.phosphosite.org/ (Table 1).

Collectively the table 1 presents current knowledge of SHARPIN phosphorylation, revealing altogether 14 phosphorylation sites from cultured cells, and five novel phosphorylation sites identified here by IVK.

### SHARPIN amino acid S146 is involved in ARP2/3 complex interaction

We selected S131 and S146 for further functional analysis due to their presence in both the IVK and *in cellulo* MS analysis (Table 1), as well as their clustering to an unstructured linker region of SHARPIN the function of which is yet unknown (Fig. 1C). To investigate the functional role of S131 and S146 phosphorylation, we created alanine mutants of these phospho-sites in a GFP-SHARPIN mammalian expression vector. Upon transient transfection, both mutants were expressed at comparable levels as WT GFP-SHARPIN when assessed by Western blotting (Fig. S1B). As functional read-outs, we used previously established assays for three SHARPIN-regulated functions: integrin activity, LUBAC activity, and ARP2/3 interaction (Bouaouina, Harburger et al., 2011, Harburger, Bouaouina et al., 2009, Khan et al., 2017).

To investigate the impact of these mutations on integrin inhibition by SHARPIN, we used the previously reported FACS assay (Bouaouina et al., 2011, Harburger et al., 2009). As expected, siRNA-mediated knock-down of SHARPIN in HeLa cells showed an increase in the integrin activity (Fig. S1C) whereas over-expression of GFP-SHARPIN-WT significantly inhibited integrin activity in these SHARPIN depleted cells (Fig. 2A). However, as both phospho-mutants also significantly inhibited integrin activity, we conclude that these phosphorylation sites are not relevant for the ability of SHARPIN to inhibit integrins (Fig 2A).

**FIG. 2.**
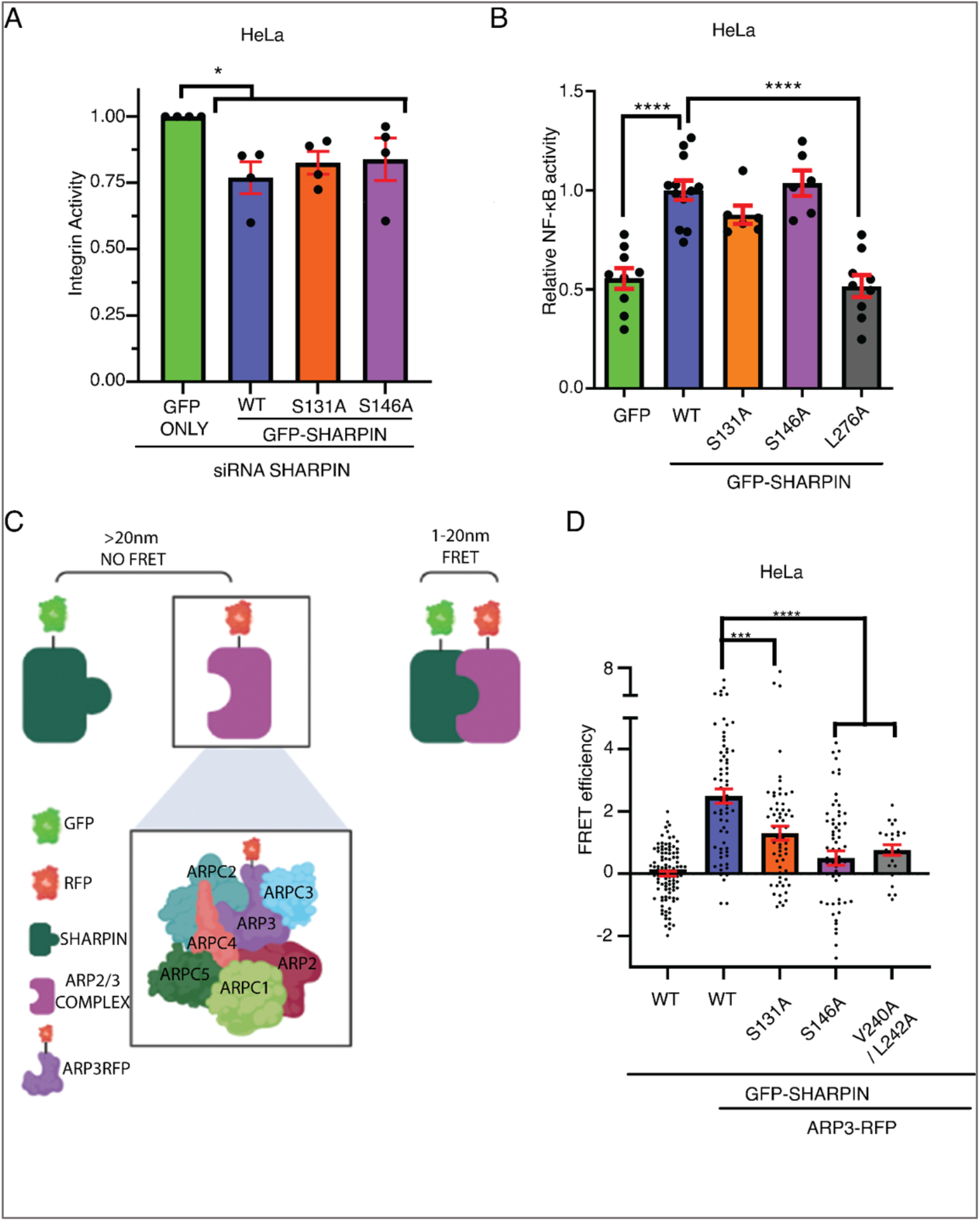
SHARPIN S131 and S146 phosphorylation promotes interaction between SHARPIN and ARP2/3 complex. (A) Quantification of integrin activity in endogenous SHARPIN silenced HeLa cells overexpressing indicated SHARPIN variants in a FACS assay (n=4). (B) TNF-induced NF-κB promoter activity in HeLa cells overexpressing indicated SHARPIN variants. NF-κB promoter activity was measured using luciferase reporter assay. GFP-SHARPIN L276A was used as a negative control (n=3). (C) Illustration of FRET between GFP-SHARPIN and ARP3-RFP. (D) Quantification of FRET efficiency in HeLa cells overexpressing indicated proteins subjected to FRET analysis by FLIM (n=5).

To analyze the effect of S131 and S146 phosphorylation sites on LUBAC activation, we used the NF-KB activity luciferase reporter assay in HeLa cells. As expected, a loss of SHARPIN significantly reduced NF-kB activity (Fig S1D), while overexpression of GFP SHARPIN-WT increased NF-kB activity (Fig. 2B). Consistent with a previous report (De Franceschi et al., 2015), the structural mutant L276A was incompetent to promote NF-kB activity (Fig. 2B). Notably, the S131A and S146A mutants were indistinguishable from SHARPIN-WT in their capacity to promote NF-kB activity (Fig. 2B), demonstrating that, like their neutral effect on integrin activity, these phosphorylation sites are not involved in regulation of LUBAC activation.

To investigate the effect of the mutations on the SHARPIN-ARP2/3 complex interaction, we analyzed Fluorescence Resonance Energy Transfer (FRET) efficiency between GFP-SHARPIN and ARP3-RFP in HeLa cells as described earlier (Fig. 2C)(Khan et al., 2017). As expected, no FRET signal was observed in cells with over-expression of GFP-SHARPIN-WT alone, whereas its co-expression with ARP3RFP resulted in a clear FRET signal (Fig. 2D). Interestingly, FRET activity in cells expressing GFP-SHARPIN S131A or S146A mutants was significantly lower as compared to the GFP-SHARPIN WT expressing cells, and the activity with S146A was indistinguishable from the structural mutant V240A/L242A used as a negative control (De Franceschi et al., 2015).

These data demonstrate that the studied SHARPIN phosphorylation sites are not involved in the integrin inhibition, or LUBAC regulation, but they significantly contribute to SHARPIN-ARP3 interaction. Out of these two mutations, S146A had clearly stronger effect on ARP3 interaction, and it was thus selected for the further functional validation.

### Constitutive SHARPIN S146 phosphorylation contributes to lamellipodium formation

ARP2/3-dependent lamellipodia formation promotes cell migration and invasion (Molinie & Gautreau, 2018, Mondal et al., 2021, Suraneni, Rubinstein et al., 2012). We have previously shown that ARP2/3 interaction with SHARPIN promotes lamellipodia formation (Khan et al., 2017). Consistent with that study, siRNA-mediated knockdown of SHARPIN in NCI-H460 lung cancer cells significantly decreased the lamellipodium formation and resulted in cells with rounded appearance (Fig. 3A). In a rescue experiment where SHARPIN-silenced cells were transfected with either GFP-only, GFP SHARPIN-WT, S146A, or V240A/L242A double mutant as a negative control (Khan et al., 2017), only the SHARPIN-WT was able to rescue the lamellipodium formation in (Fig 3B). These data are consistent with the FRET data (Fig. 2D), and indicate that phosphorylation of S146 is required for SHARPIN-mediated ARP2/3 activation.

**FIG 3.**
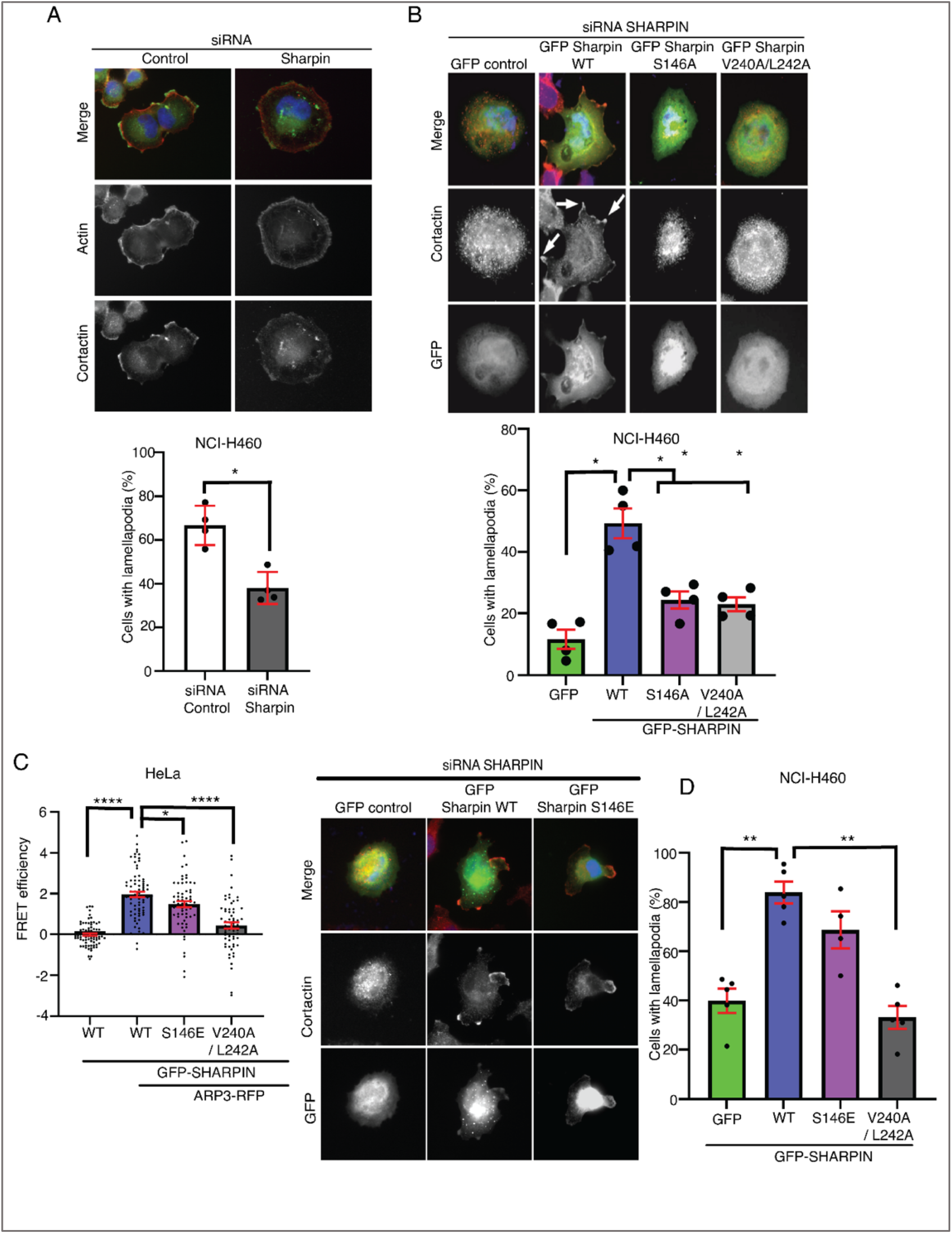
Phosphorylation of SHARPIN at S146 promotes lamellipodia formation. (A) Impact of SHARPIN silencing in NCI-H460 cell’s ability to form lamellapodia. Graph shows quantification of cells with lamellapodia (n=4). (B) Lamellipodia formation in endogenous SHARPIN silenced NCI-H460 cells overexpressing indicated Sharpin variants. GFP-SHARPIN V240A/L242A is used as a negative control. Graph shows quantification of cells with lamellapodia (n=4). (C) Quantification of FRET efficiency in HeLa cells overexpressing indicated proteins subjected to FRET analysis by FLIM (n=4). (D) Lamellipodia formation in endogenous SHARPIN silenced NCI-H460 cells overexpressing indicated Sharpin variants. GFP-SHARPIN V240A/L242A is used as a negative control. Graph shows quantification of cells with lamellapodia (n=5).

As S146 was found phosphorylated in the unperturbed cancer cells (Table I), we assumed that overexpression of phosphomimic glutamate mutant of S146 (S146E) would not impact ARP2/3 interaction or lamellipodium formation by SHARPIN. Use of the S146E mutant would also be an important control that the impaired lamellipodia formation by S146A mutant was truly due to lack of phosphorylation, and not due to structural impact of any random mutation. Importantly, although GFP-SHARPIN S146E showed slightly reduced binding to ARP3-RFP (Fig. 3C), its overexpression resulted in comparable rescue of lamellipodia formation as compared to GFP-SHARPIN WT expressing cells (Fig. 3D). Thereby we conclude that the lack of lamellipodia rescue with the S146A mutant was due to impairment of phosphorylation at S146.

### SHARPIN promotes cancer cell invasiveness

The ARP2/3 complex is a critical mediator of the entire metastatic cascade; from migration to invasion and *in vivo* metastatic spread (Molinie & Gautreau, 2018, Mondal et al., 2021). Based on the results above, we hypothesized that due to its impact on ARP2/3 complex interaction, S146 phosphorylation on SHARPIN promotes cancer cell invasiveness. This was particularly interesting as S146 phosphorylation selectively influenced lamellipodia formation without affecting the other studied signaling functions of SHARPIN (Fig. 2). To unambiguously study the function of SHARPIN in 3D invasion of cancer cells, we utilized CRISPR/Cas9-generated SHARPIN knock-out (KO) NCI-H460 lung cancer cells generated previously (Khan et al., 2017), and created additional SHARPIN KO MDA-MB-231 triple-negative breast cancer cells, and HeLa cervical cancer cells. Selection of these cell lines was due to high SHARPIN amplification frequency in these cancer types (Fig S1A). After single cell cloning of SHARPIN targeted CRISPR/Cas9 clones, Western blot analysis was used to demonstrate complete loss of endogenous SHARPIN in the MDA-MB-231 and HeLa SHARPIN KO cells (Fig S1E). Tracking the proliferation of MDA-MB-231 cell lines by Incucyte live-cell imaging for 4 days revealed no significant differences (Fig. S1F). Therefore, the potential effects of knockout of SHARPIN in 3D invasion was not confounded by significant effects on cell proliferation.

The functional contribution of SHARPIN on 3D invasion was assessed using an Inverted Transwell Invasion assay (Jacquemet et al., 2016). Remarkably, SHARPIN deletion was found essential for 3D invasion in all three cell lines (Fig. 4A-C). To rule out that this was not due to unspecific effect by the CRISPR/Cas9-mediated gene editing process, we repeated the assay with MDA-MB-231 cells from which SHARPIN was transiently knocked down by siRNA. Also in this setting, SHARPIN inhibition displayed a significant loss of invasion (Fig. 4D).

**FIG 4.**
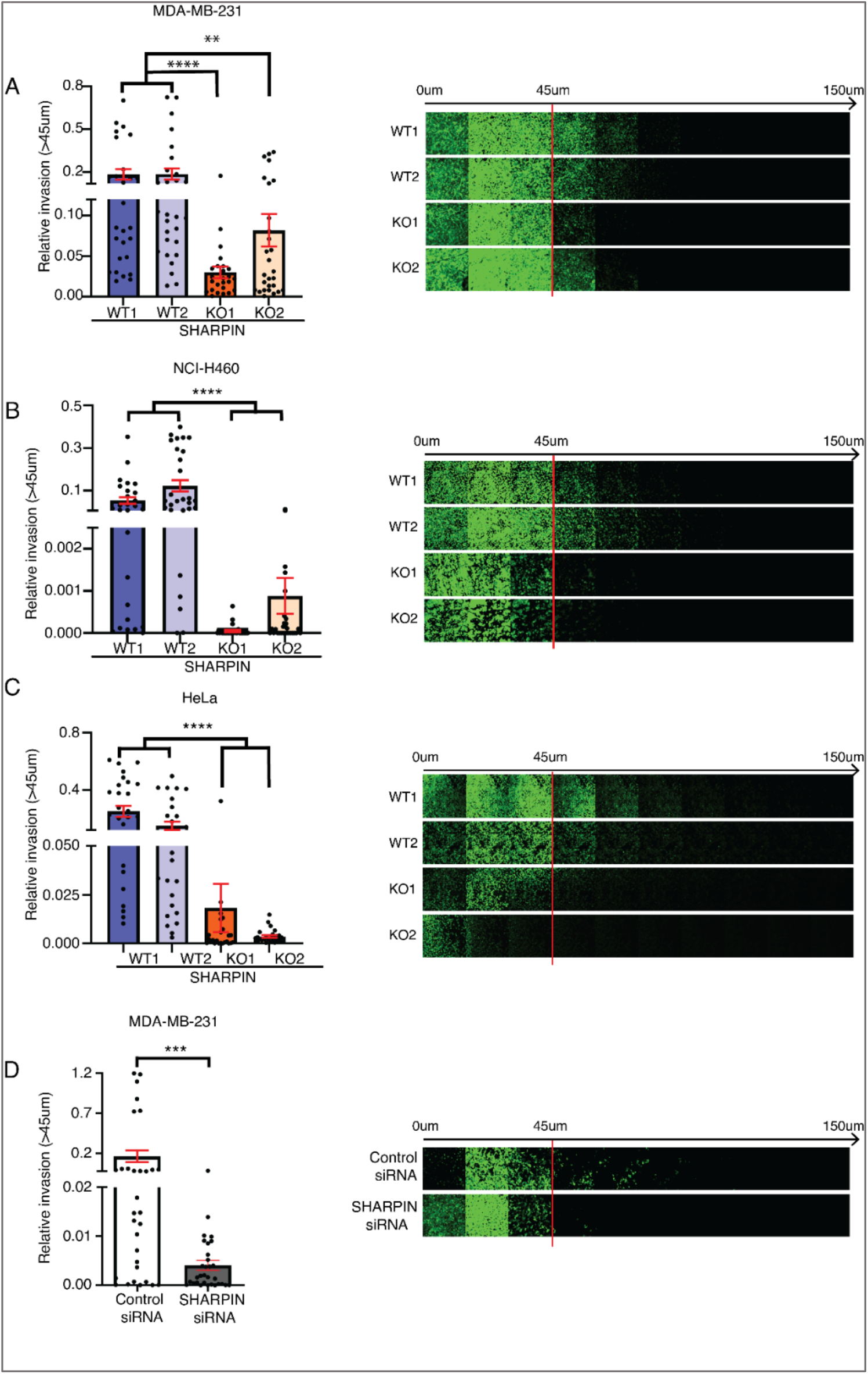
SHARPIN is essential for cancer cell invasion. (A,B,C) Impact of endogenous SHARPIN knockout on an inverted 3D invasion assay in MDA-MB-231, NCI-H460 and HeLa cells generated by CRISPR-Cas9 gene editing (n=3). (D) Relative invasion in endogenous SHARPIN siRNA silenced MDA-MB-231 cells (n=3).

These results demonstrate an essential role for SHARPIN in 3D invasion of cancer cells from three different human cancer types with high amplification frequency of SHARPIN (Fig. S1A).

### Single phosphorylation site S146 on SHARPIN determines cancer cell invasiveness

The results above demonstrate that SHARPIN S146 phosphorylation promotes lamellipodia formation (Fig. 3), which is a known requirement for cancer cells invasion, and that SHARPIN is required for 3D invasion across cancer cell lines (Fig. 4). To investigate whether S146 phosphorylation of SHARPIN can alone define the ability of cancer cells to invade, the MDA-MB-231 SHARPIN KO clones were used to generate a cell line stably expressing either GFP-only, GFP-SHARPIN-WT or GFP-SHARPIN-S146A mutant. Whereas negligible invasion was again seen with the KO cells in Inverted Transwell Invasion assay, a complete rescue was seen in cells expressing GFP-SHARPIN-WT. However, no rescue was observed in the GFP-only, or GFP-SHARPIN S146A mutant expressing cell lines (Fig. 5A). To rule out that these were clonal effects, and to expand the relevance of these findings to yet another cell model the experiment was repeated in prostate cancer PC3 SHARPIN KO cells (Fig. S1E). Consistent with the results in MDA-MB-231 cells, overexpression of GFP-SHARPIN-WT displayed significant rescue, whereas GFP-SHARPIN S146A mutant expressing cells were indistinguishable from control GFP expressing cells (Fig. 5C).

**FIG 5.**
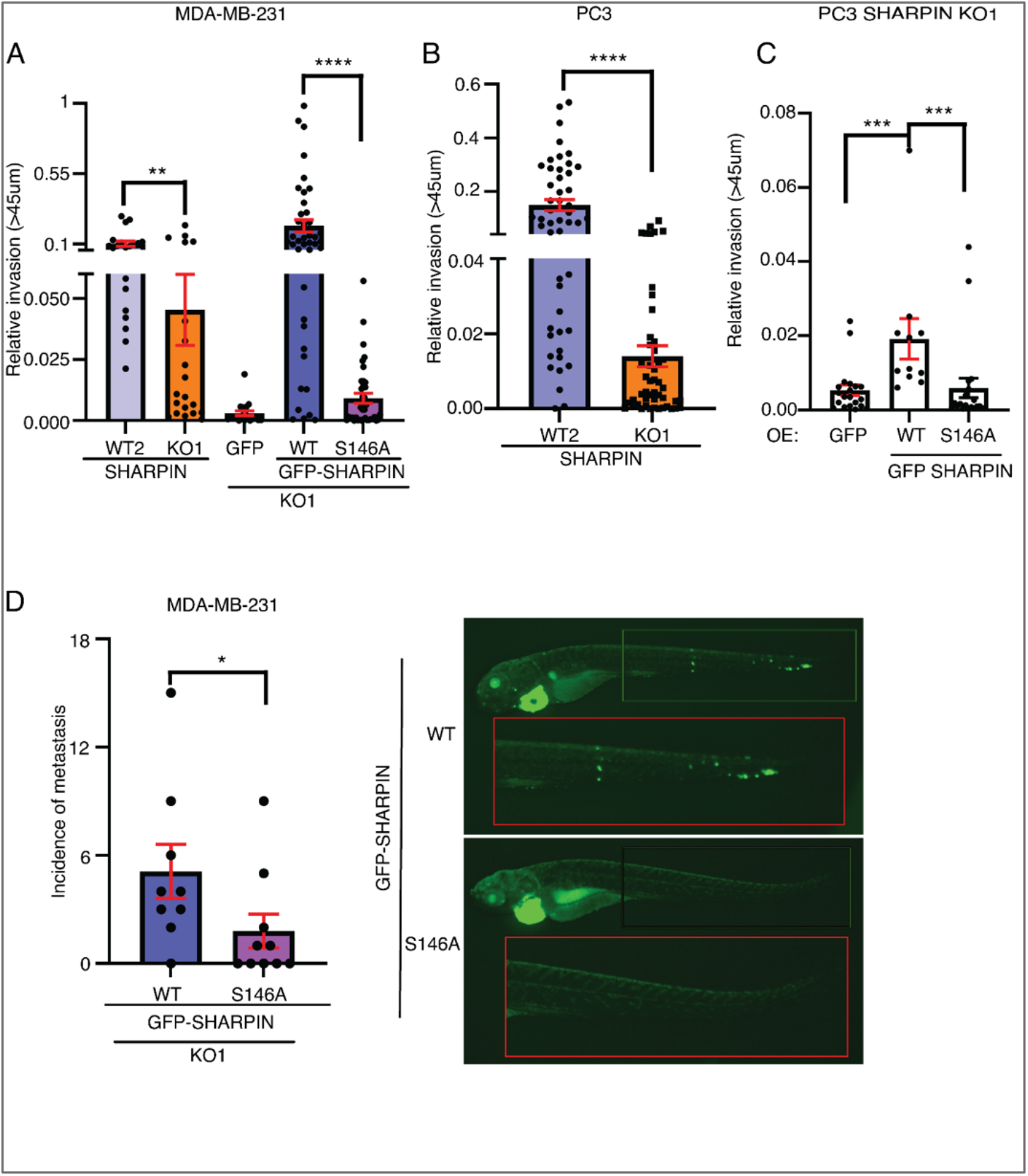
SHARPIN S146 determines cancer cell invasion and metastasis. (A) 3D invasion of MDA-MB-231 SHARPIN knock out cells stably expressing GFP only, GFP-SHARPIN WT or GFP-SHARPIN S146A (n=3). (B) 3D invasion of prostate cancer PC3 cells with knockout of endogenous SHARPIN (n=3). (C) 3D invasion of PC3 SHARPIN knock out cells overexpressing GFP only, GFP-SHARPIN WT or GFP-SHARPIN S146A (n=3). (D) In vivo zebrafish metastasis of MDA-MB 231 cells stably expressing GFP-SHARPIN WT or GFP-SHARPIN S146A mutant at day 4 following heart injection. Graph shows quantification of incidence of metastasis to zebrafish tail. (n=10).

Finally, to investigate whether these results translate into invasion phenotype in animal model, we used the zebrafish model for cancer cell invasion (Teng, Xie et al., 2013). Zebrafish embryo hearts were injected with MDA-MB-231 cells stably expressing either GFP SHARPIN-WT or the GFP-SHARPIN S146A mutant. The embryos were then fixed and imaged on day 4 following the injection. Image analysis revealed a significant decrease in distant tail metastases in embryos injected with GFP-SHARPIN S146A mutant cells as compared to the GFP-SHARPIN-WT (Fig. 5D).

Collectively these results demonstrate that SHARPIN serine 146 phosphorylation constitute a functional determinant of 3D cancer cell invasion both *in vitro* and *in vivo*.

## Discussion

Metastasis is the primary cause of cancer-related deaths in most of human solid malignancies. Thereby, identification of novel targets for anti-metastatic therapies could lead to profound decrease in cancer mortality, and increased quality of life of cancer patients(Ganesh & Massagué, 2021). In this study we demonstrate that SHARPIN is essential for 3D invasion of cancer cells from four different human cancer types, and that SHARPIN S146 phosphorylation functions as a critical invasion promoting phosphorylation switch.

SHARPIN gene amplification and protein overexpression has been observed in several human cancer types (Fig. S1A) (Bii et al., 2015, De Melo & Tang, 2015, He et al., 2010, Jung et al., 2010). SHARPIN is a multifunctional protein regulating a number of cellular pathways and functions (Gerlach et al., 2011, He et al., 2010, Ikeda et al., 2011, Landgraf et al., 2010, Lim et al., 2001, Nastase et al., 2016, Park et al., 2016, Pouwels et al., 2013, Rantala et al., 2011, Tokunaga et al., 2011), and at least some of these roles of SHARPIN are mutually exclusive (De Franceschi et al., 2015). However, it has remained a mystery how the choice between different SHARPIN functions is controlled. Here, we addressed this questions by comprehensive analysis of SHARPIN phosphorylation. By IVK assay, we demonstrated that SHARPIN is phosphorylated by major oncogenic kinases such as PKC alpha, CDK4/CycD3, ERK1 AND ERK2. On the other hand, combination of *in cellulo* phosphoproteomics analysis and database searches, we validated the amino acids that are constitutively phosphorylated in cancer cells (Table 1). Whereas previous study demonstrated the functional role of S165 phosphorylation on SHARPIN-mediated LUBAC regulation, the role of the other SHARPIN phosphorylation sites has not been studied as yet. Here we focused on functional analysis of S131 and S146 phosphorylation as these sites were observed phosphorylated both on mass spectrometry and the IVK data (Fig. 1C and Table 1).

Prior this study, SHARPIN was known to promote lamellipodium formation through interaction with the ARP2/3 complex, and it was further demonstrated that this function was independent of its LUBAC- and integrin related roles (Khan et al., 2017). Here we demonstrate role for S146 and S131 phosphorylations on SHARPIN-ARP2/3 interaction, and that mutations of these sites had no effect on the ability of SHARPIN to inhibit integrins or on NF-KB activation. S146 phosphorylation of SHARPIN was further validated to promote lamellipodia formation, but consistent with constitutive phosphorylation of S146 based on mass spectrometry data, the phosphorylation mimicking mutation (S146E) functioned as a WT. Furthermore, we demonstrate that phosphorylation of SHARPIN at S146 translates into the ability of cancer cells to invade and to metastasize *in vivo*.

In summary, our data teases out a single phosphorylation event which is essential for tumor cell invasion. Clinically this mechanism may at least partly contribute to the poor clinical outcome in patients with high SHARPIN expression. Therefore future studies should be directed to validate S146 phosphorylation in patient samples in correlation with patient metastasis status. Related to development of future anti-metastatic therapies, our data provide very convincing evidence that inhibition of SHARPIN expression effectively abrogates 3D invasion across cells from different cancer types. Further, future structural analysis of ARP2/3 bound to SHARPIN unstructured region between the PH and UBL domains could reveal important clues for potential targetability of this cancer cell invasion promoting protein-protein interaction.

## AUTHOR CONTRIBUTIONS

**U.B**: Designing experiments, Conducting Experiments, Acquiring Data, Analyzing Data, statistical Analysis, Writing the manuscript.

**M.H.K**: Sharpin Knockout cell line generation

**J.P**: Designing experiments, IVK assay

**JW**: Designing experiments, writing the Manuscript

## ACKNOWLEDGEMENTS

The authors would like to thank Taina Kalevo-Mattila for technical assistance and the entire Turku Bioscience personnel for excellent working environment. We acknowledge Cell Imaging and Cytometry core facility, Proteomics core facility and Zebrafish core facility at Turku Bioscience Center supported by Biocenter Finland. The project was funded by Academy of Finland (to J.P), Finnish Cancer Foundation (to J.W) and Sigrid Juselius Foundation (to J.W). U. Butt was supported by Finnish Cultural foundation and Albin Johanssons foundation.

